# Effects of multiple sources of genetic drift on pathogen variation within hosts

**DOI:** 10.1101/190918

**Authors:** David A. Kennedy, Greg Dwyer

## Abstract

Changes in pathogen genetic variation within hosts alter the severity and spread of infectious diseases, with important implications for clinical disease and public health. Genetic drift may play a strong role in shaping pathogen variation, but analyses of drift in pathogens have oversimplified pathogen population dynamics, either by considering dynamics only at a single scale (within hosts, between hosts), or by making drastic simplifying assumptions (host immune systems can be ignored, transmission bottlenecks are complete). Moreover, previous studies used genetic data to infer the strength of genetic drift, whereas we test whether the genetic drift imposed by pathogen population processes can be used to explain genetic data. We first constructed and parameterized a mathematical model of gypsy moth baculovirus dynamics that allows genetic drift to act within and between hosts. We then quantified the genome-wide diversity of baculovirus populations within each of 143 field-collected gypsy moth larvae using Illumina sequencing. Finally, we determined whether the genetic drift imposed by host-pathogen population dynamics in our model explains the levels of pathogen diversity in our data. We found that when the model allows drift to act at multiple scales, including within hosts, between hosts, and between years, it can accurately reproduce the data, but when the effects of drift are simplified by neglecting transmission bottlenecks and stochastic variation in virus replication within hosts, the model fails. A *de novo* mutation model and a purifying selection model similarly fail to explain the data. Our results show that genetic drift can play a strong role in determining pathogen variation, and that mathematical models that account for pathogen population growth at multiple scales of biological organization can be used to explain this variation.

## Introduction

Pathogen genetic variation can have important consequences for human health, in both clinical and epidemiological settings (Alizon et al. 2011). In particular, high variation within hosts can lead to severe disease symptoms within individuals, and rapid disease transmission within populations (Read and Taylor 2001; Vignuzzi et al. 2006). An understanding of the mechanisms determining pathogen variation might therefore lead to novel interventions, reducing the toll of infectious diseases. Development of such an understanding requires quantification of the effects of population processes on pathogen genetic variation, in turn requiring mathematical models that relate population processes to genetic change.

Such models, however, tend to greatly simplify pathogen biology. Selection-mutation models, for example, often assume that pathogen populations are effectively infinite (Lorenzo-Redondo et al. 2016). Models that allow pathogen population sizes to be finite typically neglect pathogen population processes either within hosts in acute infections (Koelle et al. 2006), or between hosts in chronic infections (Pennings et al. 2014). Models that attempt to capture both of these scales of disease dynamics have assumed either that pathogen population growth within hosts is very simple (Klinkenberg et al. 2017), or that pathogen bottlenecks at transmission are complete (Didelot et al. 2014; Klinkenberg et al. 2017; Ypma et al. 2013), so that every infection begins as a clonal lineage. These simplifications could strongly alter conclusions about the effects of genetic drift on pathogen diversity, and indeed have been highlighted as key challenges in phylodynamic inference (Frost et al. 2015).

Genetic drift is a change in an allele’s frequency due to the chance events that befall individuals. The effects of drift are thus strongest in small populations, in which a few events can have a large impact (Nagylaki 1992). The high population sizes typical of severe infections have led some authors to argue that drift has little effect on pathogens (Kouyos et al. 2006; Maldarelli et al. 2013), but pathogen population sizes typically fluctuate by several orders of magnitude over the course of infection and transmission. It therefore seems likely that, when pathogen populations are small and variable, pathogen genetic variation will be strongly affected by drift. Indeed, analyses that allow for finite population sizes have shown that drift has at least weak effects on some pathogens (Pennings et al. 2014).

New infections are typically initiated by small pathogen population sizes within hosts (Gutiérrez et al. 2012), leading to bottlenecks at the time of transmission that may drive genetic drift. Pathogen population sizes within hosts can also remain small for long periods following exposure (Kennedy et al. 2014). In small populations, chance events such as the timing of reproduction can strongly influence population growth, a phenomenon known as “demographic stochasticity” (Kot 2001). When the effects of demographic stochasticity are strong, chance may allow some virus strains to replicate and survive while others go extinct, providing a second source of genetic drift that we refer to as “replicative drift”. Note that we use the term “strain” to mean a population of pathogen particles that have identical genetic sequences.

Many previous studies of genetic drift in pathogens have focused only on population processes that operate within hosts, either during experiments with model organisms, or during the treatment of human patients (Abel et al. 2015; Gutiérrez et al. 2012). Pathogen variation in nature, however, is also affected by processes that operate at the host population level, such as fluctuating infection rates during epidemics (Grenfell et al. 2004). Studies of Ebola (Azarian et al. 2015) and tuberculosis (Lee et al. 2015), for example, have shown that much of the variation present at the population level often cannot be explained by natural selection, and must instead be due to neutral processes that presumably include genetic drift.

Genetic drift in pathogens may thus be driven by population processes at multiple scales. These multiple scales can be incorporated into a single framework by constructing “nested” models, in which sub-models of within-host pathogen population growth are nested in models of between-host pathogen transmission (Mideo et al. 2008). The computing resources necessary to analyze such complex models have only become available recently, however, and so it has not been clear whether sufficient data exist to test nested models of drift (Gog et al. 2015). Indeed, even for models that assume that the strength of drift is constant across hosts, robust tests of the model predictions require both genetic data and mechanistic epidemiological models (Didelot et al. 2014), a combination that is rarely available. Whether nested models can be of practical use for understanding pathogen genetic variation in nature is therefore unclear.

For baculovirus diseases of insects, pathogen population processes have been intensively studied at both the host population level (Elderd 2013), and at the individual host level (Kennedy et al. 2014). Baculoviruses cause severe epizootics (= epidemics in animals) in many insects (Moreau and Lucarotti 2007), including economically important pest species such as the gypsy moth (*Lymantria dispar*) that we study here (Woods and Elkinton 1987). Collection and rearing protocols for the gypsy moth have long been standardized (Elkinton and Liebhold 1990), and so previous studies of the gypsy moth baculovirus *Lymantria dispar* multiple nucleopolyhedrovirus (LdMNPV) have produced parameter estimates for both within-host (Kennedy et al. 2015) and between-host (Elderd et al. 2008, 2013; Fuller et al. 2012) models. Moreover, collection of large numbers of virus-infected individuals is straightforward (Woods and Elkinton 1987), making it possible to use high-throughput sequencing methods to characterize pathogen diversity across many virus-infected hosts. Here we use a combination of whole-genome sequencing and parameterized, nested models to quantify the effects of genetic drift on the gypsy moth baculovirus. We show that a mechanistic model of genetic drift can explain variation in this pathogen, but only if the model takes into account the effects of drift at multiple scales of biological organization.

## Results

Sequencing the virus populations from each of 143 field-collected insects showed that there is substantial genetic variation in baculovirus populations between hosts. We generated consensus sequences for each of our 143 samples (see Supplemental Information A), and comparisons between consensus sequences identified 712 segregating sites at the between host scale (defined as sites where alternative variants were the consensus in more than 6 samples (≈ 5%)). These sites correspond to approximately 0.4% of the genome. Analysis of the variation at these 712 sites within each sampled virus population showed that these sites were polymorphic in some hosts but not others, which might occur if some hosts were exposed to multiple strains of virus, while others were exposed to only a single strain. We summarize genetic variation within hosts using mean nucleotide diversity (Nei and Li 1979), the probability that two randomly selected alleles at a segregating site are different (Supplemental Information A). Our conclusions were nevertheless unchanged when we used alternative metrics of diversity, such as the proportion of polymorphic loci, the effective number of alleles, or the relative nucleotide diversity (Supplemental Information I).

Measured across the consensus sequences of our 143 samples, nucleotide diversity at our 712 segregating sites was quite high at 0.404. Within samples, nucleotide diversity at these same sites ranged from 0.002 to 0.284 (mean = 0.072, s.d. = 0.077, Supplemental Information B). Overall nucleotide diversity within samples ranged from 0.001 to 0.003, with a mean of 0.001. In Supplemental Information B, we show that these values imply that a large fraction of nucleotide diversity within hosts can be explained by just 712 segregating sites, or 0.4% of the genome.

Together, these patterns suggest that substantial pathogen diversity within hosts is likely acquired from the exposure of host insects to multiple virus strains. If diversity had instead been generated by *de novo* mutation, nucleotide diversity between samples would have been less variable (Supplemental Information E), and polymorphism would have likely been spread across many sites, including sites that were not polymorphic at the population level. Immune-system mediated diversifying selection is also an unlikely explanation, because insects lack clonal immune cell expansion (Vilmos and Kurucz 1998), because immune cell expansion does not explain why some hosts have substantially more pathogen diversity than others, and because we found no evidence of diversifying selection in our sequence data (Supplemental Information H). Negative correlations between host families in susceptibility to different pathogen genotypes constitute yet a third unlikely explanation, because in the gypsy moth such correlations are positive (Hudson et al. 2016). Migration of virus or infected larvae from nearby locations with different virus strains similarly cannot explain the data, because population structure in the gypsy moth virus is minimal (Fujita 2007, Supplemental Information A).

Genetic drift, however, can explain the data, but only if we allow for effects of population processes at multiple scales of biological organization. To explain why, we first use a nested model of pathogen population dynamics (fig. 1) to show how genetic drift in pathogen populations may operate at three scales; within hosts, within epizootics, and between years. We then show that the model can only explain the data if it includes effects of drift both during transmission bottlenecks and virus growth within hosts.

**Figure 1:**
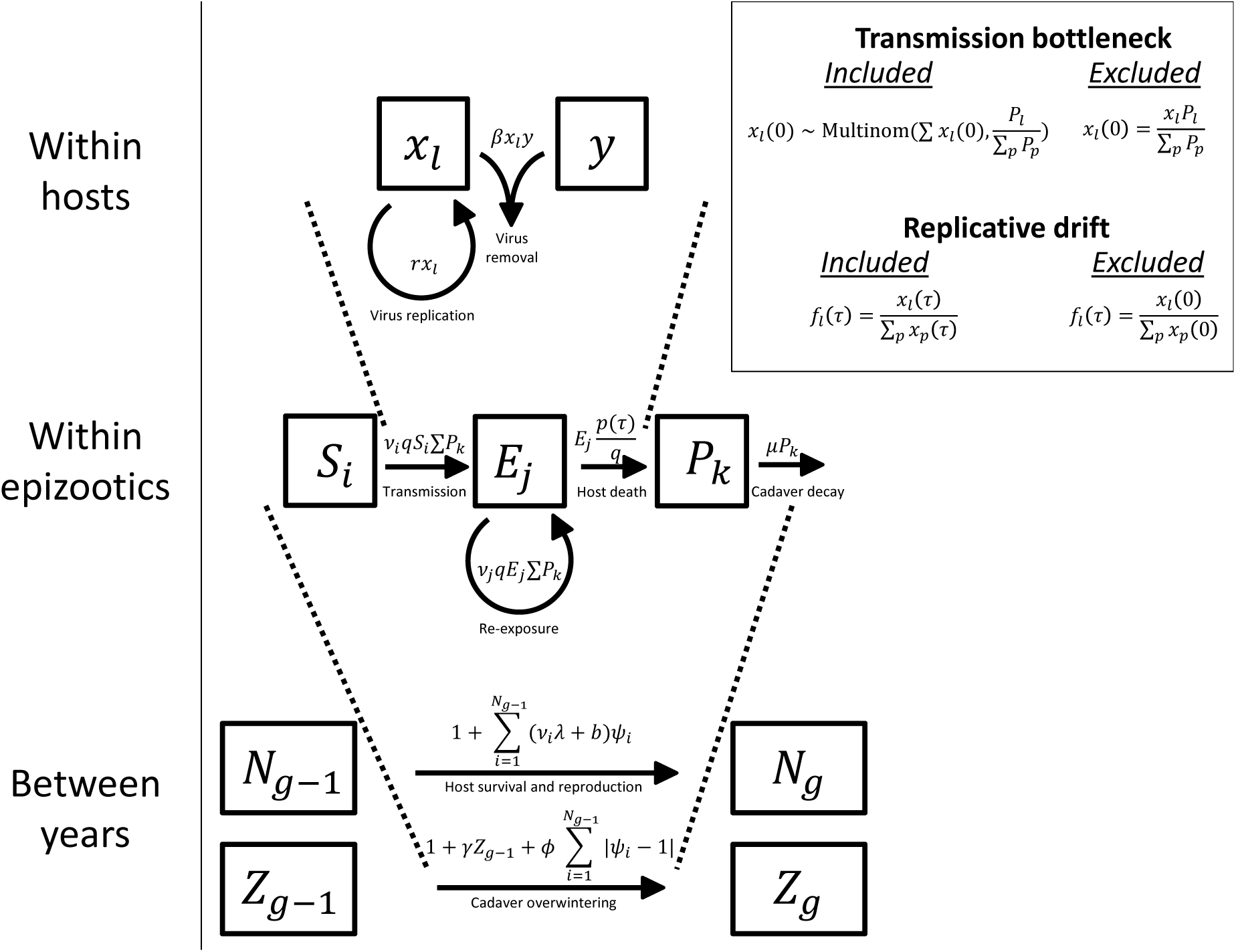
Schematic of the nested model. Bottom, the host population size *N*_*g*_ and the infectious cadaver population size *Z*_*g*_ in generation *g* depend on host and pathogen population sizes in generation *g* — 1 and the disease dynamics in that generation. Following the epizootic, surviving hosts reproduce and virus-killed cadavers overwinter at rate *φ* to start the epizootic in the following year. Middle, the disease dynamics in generation *g* — 1 follow a stochastic SEIR model (Keeling and Rohani 2008), such that a susceptible host *S*_*i*_ becomes exposed *E*_*j*_ to infectious cadaver *P*_*k*_ at rate *v*_*i*_*q*, where *v*_*i*_ is the risk of exposure for host *i* and *q* is the probability of death given exposure, which arises from the within host virus dynamics. Note that the “Removed” class *R*, corresponding to inactivated cadavers, is not explicitly shown. The probability of a host dying from virus infection at time *τ* post exposure *p*(*τ*), is determined by the dynamics of the pathogen within a host. *q* is related to *p*(*τ*) in that 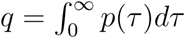. Top, within a host, virus particles *x* can reproduce or interact with immune cells *y*, resulting in the removal of both the virus particle and the immune cell. An infection fails to kill the host if all virus particles are cleared so that *x* = 0, but the host dies if the total number of virus particles reaches an upper threshold *C*. Further details are in Supplemental Information C. To produce a model that lacks replicative drift, we assume that the frequency of a virus strain *l* at time of death *τ*, *f*_*l*_(*τ*), is equal to the frequency of that strain immediately after the time of exposure *f*_*l*_(0). To produce a model that lacks transmission bottlenecks, we assume that the number of copies of a virus strain *l* at the beginning of an infection *x*_*l*_(0) is equal to the total number of virus particles that invade the host ∑_*l*_ *x*_*l*_(0), times the relative frequency of that virus strain in the cadaver that caused exposure *P*_*l*_/(∑_*p*_ *P*_*p*_). In the model that lacks transmission bottlenecks and replicative drift, host death occurs only if a larva was susceptible to one or more of the virus strains in the cadaver to which it was exposed. If so, the virus strains that the host was susceptible to are released upon host death at frequencies equal to those in the infecting cadaver.

Simulations of our within-host model show that the combination of transmission bottlenecks and replicative drift can substantially reduce pathogen diversity within hosts (fig. 2A-C). Demographic stochasticity, which is manifest in the figure as jaggedness in the model trajectories, is strongest shortly after exposure, when the pathogen population size is small. This stochasticity generates variability in the time to host death, and it also drives replicative drift. Comparing this model to a linear birth-death model (Supplemental Information C) shows that the immune system substantially slows the growth of the virus population early in the infection, which strengthens the effects of replicative drift.

**Figure 2:**
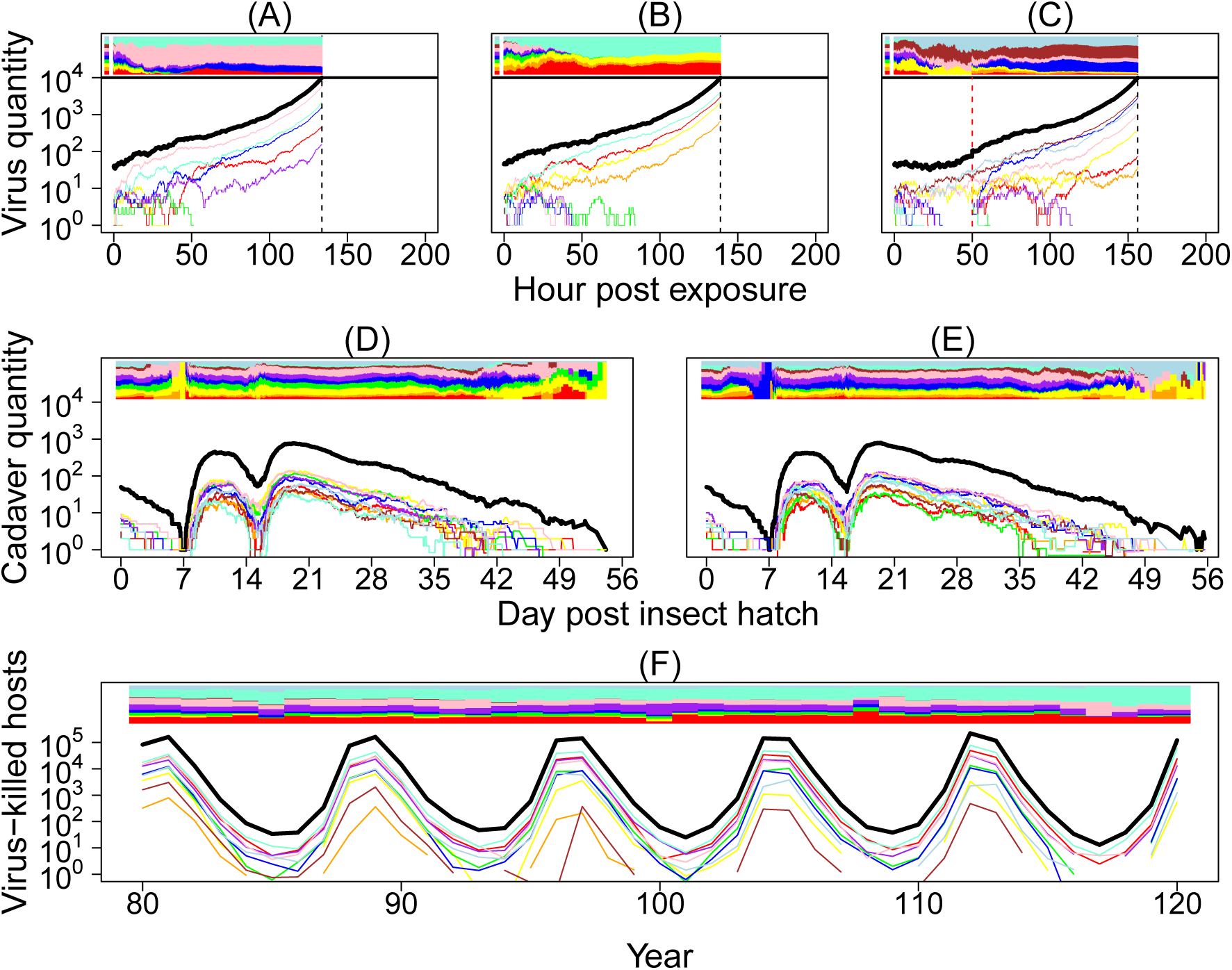
Simulations of the nested model. In all panels (A)-(F), colored curves represent the pathogen population sizes of different virus strains, and the black curve shows the total pathogen population size. The colored bar at the top of each panel shows the relative frequencies of virus strains over time. Panels (A)-(C) show three realizations of the within-host virus growth model. A re-exposure event, marked by a dashed, vertical red line, is also shown in panel (C). The top colored panel left of time 0 shows the frequency of virus strains in a cadaver that a host was exposed to at time 0 (and re-exposed to at time 50 in panel (C)). Death occurs when the total number of virus particles within a host hits an upper threshold. To aid visualization, here we set the pathogen population size at host death to be 10^4^, as opposed to the more realistic value of 10^9^ that we use when comparing our models to data. The time of death differs between simulations due to demographic stochasticity in virus growth, and in each simulation it is marked by a dashed, vertical black line. Panels (D) and (E) show two realizations of our stochastic SEIR-type epizootic model starting from identical initial conditions. Note that the curves here show cadaver quantities, rather than virus particles as in panels (A)-(C). Epizootics are initiated by overwintered cadavers that infect emerging larvae. As these cadavers decay, total cadaver quantity drops to low levels, such that the pathogen population is almost entirely composed of virus particles inside living hosts. These hosts then die initiating future rounds of infections. Panel (F) shows a realization of our pathogen model, with trajectories showing the total number of virus-killed hosts in each generation. The frequency of pathogen strains can drift over time, an effect that is particularly noticeable during troughs of infection.

Overwintered virus infects hatchlings during the initial emergence of hosts from eggs, an effect that is apparent in our simulations of the epizootic model (fig. 2D-E). After the overwintered virus decays, there is a short period when cadavers are rare, such that the vast majority of virus is present only within exposed larvae. When these exposed larvae die, the virus that they release is transmitted to new larvae feeding on foliage. During this time, the relative frequencies of different virus strains consumed by larvae can fluctuate strongly due to the drift that occurs when cadavers are rare. Low densities of cadavers can thus alter the relative frequency of strains within hosts. In the figure, the initial host population consists of more than 10,000 hosts, reflecting the high densities at which baculovirus epizootics occur in insect populations in nature (Moreau and Lucarotti 2007). Demographic stochasticity nevertheless influences the composition of virus strains near the end of the epizootic, when the pathogen population begins to die out, in turn allowing drift to influence which virus strains cause infections within hosts.

Over longer time periods, fluctuations at the population scale (fig. 2F) produce host-pathogen cycles that match the dynamics of gypsy moth outbreaks in nature (Dwyer et al. 2000; Elderd et al. 2008). These large fluctuations can drive changes in the relative frequency of pathogen strains, especially when pathogen population sizes and overall infection rates are at their lowest, in the troughs between host population peaks. Host-pathogen population cycles in our model thus further strengthen the effects of genetic drift on the pathogen.

Our combined model therefore shows that drift can act both within and between hosts, and at time scales ranging from hours to decades. To test the model, we compared its predictions of nucleotide diversity to the levels of nucleotide diversity in our data. To test whether the data can be explained equally well by models that neglect one or more sources of drift, we also tested models that eliminated replicative drift, or that eliminated both replicative drift and transmission bottlenecks. Note that it is not possible to construct a model that includes replicative drift but not transmission bottlenecks, because replicative drift requires virus population sizes to be integer values, and forcing the virus population to have an integer value necessarily imposes a form of bottleneck. Also, to test whether the data are better explained by selection than by drift, we constructed a model that allows for purifying selection to act within hosts, but that lacks both replicative drift and transmission bottlenecks.

These comparisons show that only the model that includes both replicative drift and bottlenecks can explain the data (fig. 3). The neutral model that includes only drift at the host-population scale predicts within-host diversity levels that are much higher and much less variable than in the data. The model that includes population-scale drift and bottlenecks but not replicative drift, and the model that includes purifying selection but not transmission bottlenecks or replicative drift both correctly predict that there will be substantial variation across hosts, but they predict diversity levels that are much higher than in the data. In Supplemental Information D and G, we show the that these qualitative conclusions are robust across parameter values that determine bottleneck severity and selection intensity. In contrast, the model that includes replicative drift and transmission bottlenecks accurately predicts the entire distribution of diversity levels seen in the data. This visual impression is strongly confirmed by differences in the Monte Carlo estimates of the likelihood scores across models (Supplemental Information F: neutral model with neither bottlenecks nor replicative drift, median log mean likelihood = −503.2; purifying selection model, median log mean likelihood = −353.1, neutral model with bottlenecks but not replicative drift, median log mean likelihood = −266.7; neutral model with both bottlenecks and replicative drift, median log mean likelihood = −63.9). Because no parameters were fit to the diversity data, we do not need a model complexity penalty, but the difference in the number of parameters across models was in any case dwarfed by the differences in the likelihood scores.

**Figure 3:**
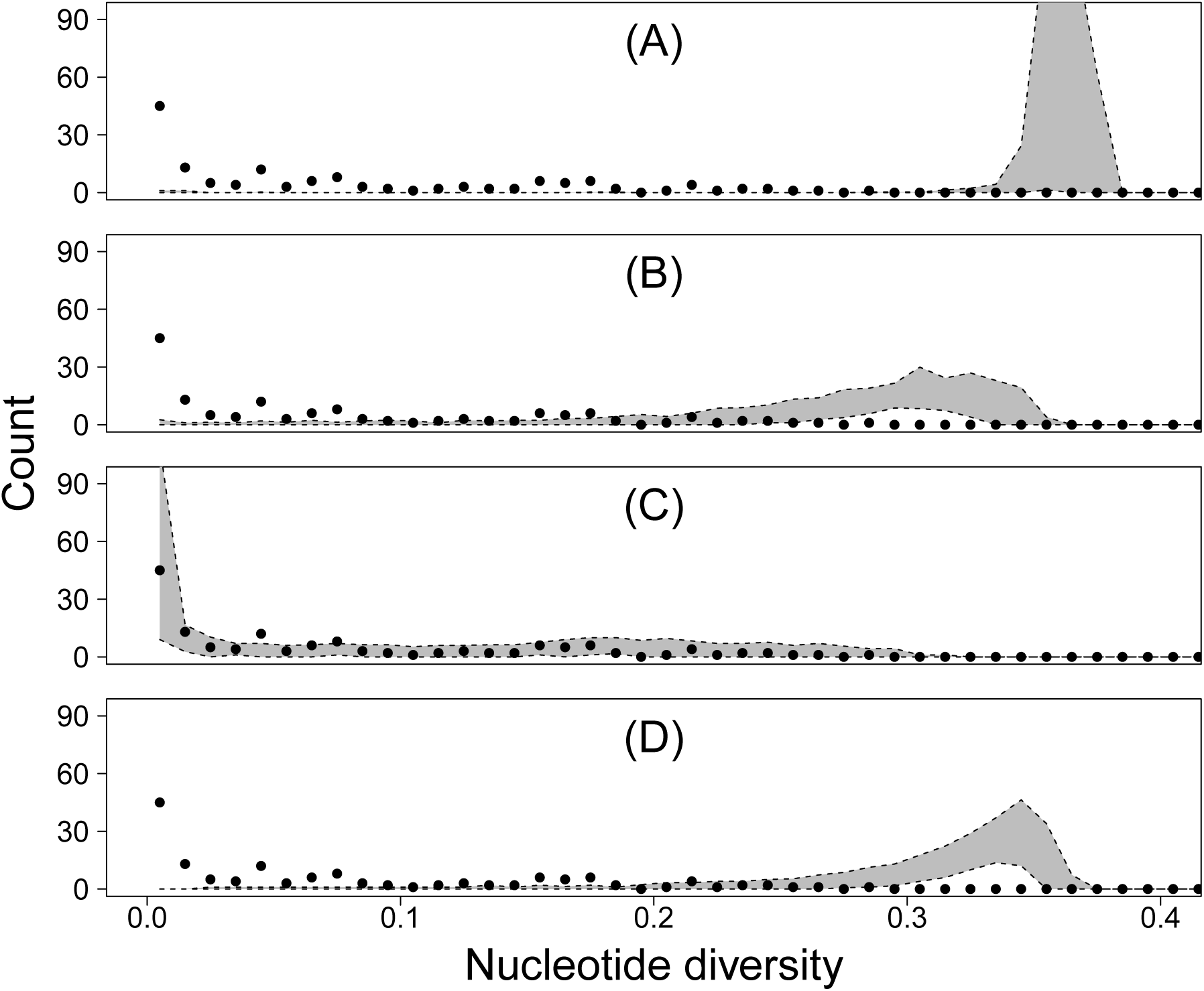
Comparison of the predictions of our models (gray-shaded areas, showing 95 percent confidence intervals of model realizations) to the distribution of nucleotide diversity within 143 individual infected hosts calculated from our sequence data (black dots show data on nucleotide diversity within hosts). (A) shows the predictions of a model that lacks both transmission bottlenecks and replicative drift, (B) shows the predictions of a model that includes transmission bottlenecks but not replicative drift, (C) shows the predictions of a model that includes both transmission bottlenecks and replicative drift, and (D) shows the predictions of a model that includes purifying selection within hosts but not transmission bottlenecks or replicative drift.

Our results thus show that a model that accounts for the effects of population processes at multiple scales can explain differences in pathogen variation across hosts in the gypsy moth baculovirus. In contrast, models that simplify the effects of genetic drift by ignoring effects of transmission bottlenecks and replicative drift, or that allow for selection but not within host drift, cannot explain the diversity of this pathogen. More broadly, because the model parameters were estimated entirely from experimental data on baculovirus infection rates (Supplemental Information C), we are effectively carrying out cross-validation of the model.

The highly skewed distribution of nucleotide diversity apparent in our data can thus be explained by a model that allows for drift at multiple scales, and that includes multiple sources of drift within hosts, but not by simpler models. In addition, fig. 4 shows that the best model can reproduce entire distributions of diversity within individual hosts. Allele-frequency distributions in the model nevertheless tend to have slightly shorter tails and narrower peaks compared to the data. These mild discrepancies may be partially explained by mutations that occurred during viral passaging or during library preparation, but they can also be explained by small biases introduced during the mapping of our short sequence reads to the reference genome (Supplemental Information J). The data therefore do not reject the model.

**Figure 4:**
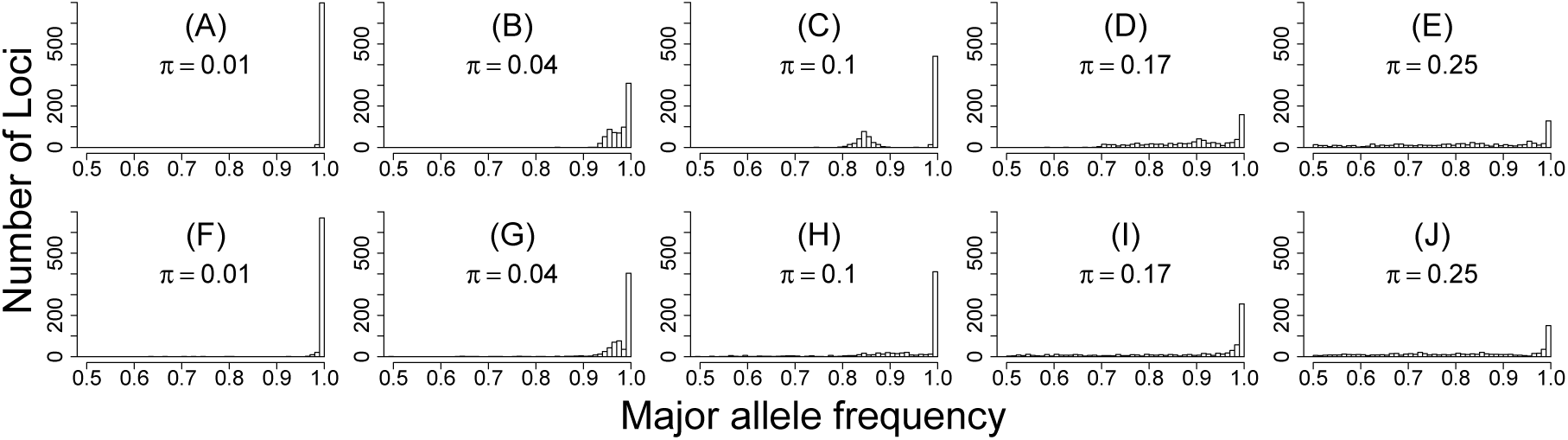
Representative distributions of allele frequencies from individual hosts in our best model (A-E) and in our data (F-J). Each plot shows the distribution of allele frequencies within a single individual at 712 segregating sites, showing only the frequency of the most common allelic variant at each locus within that host. The number on each plot is the mean nucleotide diversity within that particular host. Model plots are aligned with similar data plots. The lack of diversity in panels A and F suggests that the virus population within these hosts consist of only a single virus strain. The bimodal distributions in panels B, C and G suggest that these virus populations contain exactly two virus strains. The high diversity but lack of bimodality in panels D, E, H, I, and J suggests that these virus populations consist of more than two virus strains.

Our virus samples were collected at times of peak or near-peak gypsy moth densities, which are the only times at which large numbers of larvae can be collected easily, and so the data do not directly show how changes in pathogen population size at the host-population scale affect pathogen variation. We therefore used our best model to explore how pathogen variation within hosts will change over the course of the gypsy moth outbreak cycle. Within-host diversity is predicted to be highest just as the host population begins to crash due to the pathogen, after which diversity is predicted to gradually decline until the next outbreak (fig. 5). Reductions in within-host variation due to transmission bottlenecks and replicative drift are thus counter-balanced by increases in within-host variation at the time of host population peaks, due to the high frequency of multiple exposures when host populations are large. Long-term, population-scale processes can therefore also strongly affect within-host variation.

**Figure 5:**
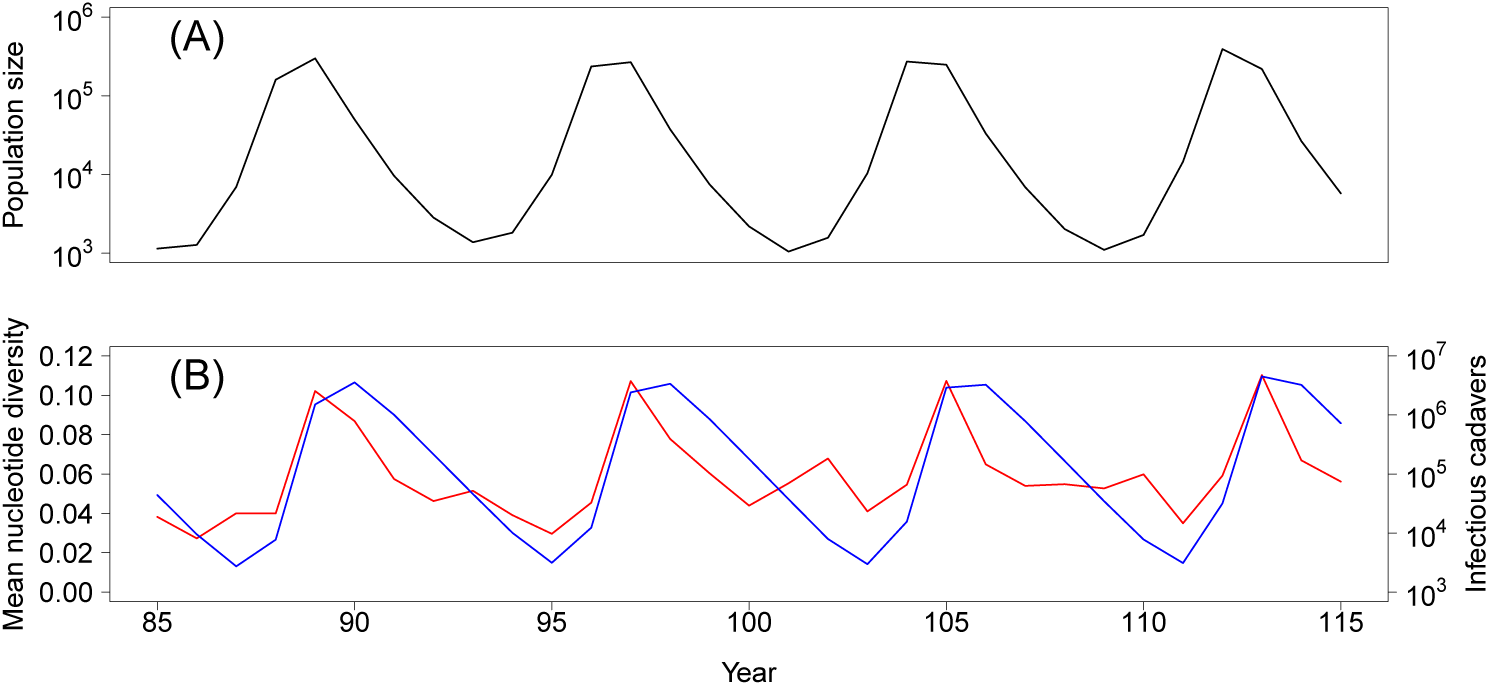
Model predictions of the effects of changes in the populations of susceptible and infected hosts on within-host pathogen diversity, over the host-pathogen population cycle. (A) The population size of uninfected hosts. (B) The population size of infectious cadavers (blue) and the mean nucleotide diversity (red).

## Discussion

A basic prediction of population genetics theory (Nagylaki 1992), and a fundamental assumption of phylodynamic modeling (Grenfell et al. 2004), is that the effects of genetic drift are determined by population processes. Explicit tests of this assumption for infectious diseases, however, are rare. We used a model that was developed and parameterized using non-genetic datasets to show that patterns of genetic diversity in an insect pathogen can be explained by a model that accounts for population processes at multiple scales, but not by models that simplify or neglect the effects of drift. Previous work has attempted to infer disease demography and pathogen evolution from genetic data (Grenfell et al. 2004). Here we instead began with an existing population process model that has already been fit to epidemiological data, and we use it to predict pathogen genetic data. We thus tested the extent to which disease demography can be used to predict neutral pathogen evolution.

A simple model of purifying selection was not able to explain the patterns of diversity in our data. Models that instead invoke diversifying selection or more complex patterns of host-specific immune selection might provide reasonable explanations for our data, but such models require extra parameters to account for the costs and benefits of alternative alleles, increasing the complexity of the models (Orr 1998). Moreover, drift is an inherent property of small populations, and so models that invoke selection should still allow for effects of drift if population sizes are small. In our case, complex models of selection were not needed to explain patterns of diversity, suggesting that the effects of selection on our data are weak relative to the effects of drift. Selection may nevertheless be necessary to explain variation in other pathogens or other datasets. Given that polymorphism has been widely observed in insect baculoviruses (Chateigner et al. 2015; Hodgson et al. 2001), our results suggest that baculoviruses present opportunities to understand the relationship between host-pathogen ecology and pathogen diversity.

We have shown that both transmission bottlenecks and replicative drift have detectable impacts on pathogen diversity. Due to the difficulty of separating these effects, previous studies of genetic drift have assumed that bottlenecks are complete (Klinkenberg et al. 2017; Pennings et al. 2014; Ypma et al. 2013), have ignored impacts of key biological processes such as the host immune response (Sobel Leonard et al. 2017), or have summarized the effects of multiple sources of drift with a single parameter, the effective population size *N*_*e*_ (Volz et al. 2017). Similarly, estimates of bottleneck size often combine the effects of transmission bottlenecks and replicative drift into a single estimate of the effective bottleneck, biasing estimates of transmission botteleneck size (Sobel Leonard et al. 2017, Supplemental Information D). Distinguishing between transmission bottlenecks and replicative drift, however, may provide novel insights into disease control strategies. For example, the emergence of resistance to antibiotic drugs might be slowed if drug therapy windows are restricted to periods when the effects of replicative drift are strongest, such as when pathogen populations are small or are turning over rapidly.

To show that both transmission bottlenecks and replicative drift play an important role in shaping pathogen diversity within hosts, we have focused our analysis on common variants that cannot be easily explained by *de novo* mutation. Additional variation is nevertheless present (Supplemental Information B). In our case, this other variation occurs at such low levels that it cannot be readily distinguish from sequencing error, but it is almost certainly true that mutation and selection also play roles in shaping total pathogen diversity within hosts. Our argument is therefore not that mutation and selection are unimportant, but instead that transmission bottlenecks and replicative drift can strongly affect pathogen diversity within hosts. In our case, bottlenecks and replicative drift appear to be the main drivers of diversity at sites that segregate at the population level.

High-throughput sequencing has revolutionized our ability to measure pathogen variation. It has been used to detect drug-resistance (Mideo et al. 2016), to discover novel viruses in nature (Lipkin and Anthony 2015), and to diagnose disease in clinical settings (Wilson et al. 2014). Our work shows that high throughput sequencing can also provide important insights into the ecology and evolution of host-pathogen interactions, especially when combined with nested disease models. The increasing availability of both parameterized models (Keeling and Rohani 2008) and genomic data (Hatherell et al. 2016) suggests that our approach of using genetic data to challenge models of nested disease dynamics may be widely applicable.

## Methods

### Model description

The gypsy moth baculovirus LdMNPV, is a double stranded DNA virus belonging to the family *Baculoviridae*. The virus is approximately 161 kb, and like all baculoviruses, it exists in two forms, as an “occlusion body” that is highly stable in the environment due to its protective proteinaceous matrix, and as a “budded virus” that is released from cells during replication within hosts.

The gypsy moth baculovirus is transmitted when larvae consume occlusion bodies while feeding on foliage (Elderd et al. 2008). If the resulting virus population grows inside the host to a sufficiently large size, the larva dies, releasing new occlusion bodies onto the foliage. These occlusion bodies are then available to be consumed by additional conspecifics (the virus is species specific (Moreau and Lucarotti 2007)), leading to very high infection rates in high density populations (Woods and Elkinton 1987). During the fall and winter, when the insect is in the egg stage, the virus persists beneath egg masses laid on cadavers or other locations where the virus may be protected from degradation by ultraviolet light (Fleming-Davies and Dwyer 2015; Murray and Elkinton 1989). Genetic drift in the gypsy moth baculovirus may therefore be affected by population processes at multiple scales, including within individual hosts and across the host population.

Exposure to the virus results in an initial population of only a few virus particles (Kennedy et al. 2014; Zwart et al. 2009), and the population size in the host remains small for a substantial period of time following exposure (Kennedy et al. 2014, 2015). Our model of pathogen growth within hosts therefore tracks population sizes from the initial population bottleneck through the stochastic growth of the pathogen population, until death or recovery. Our model thus explicitly includes genetic drift (fig. 1).

Our within-host model is based on a birth-death model (Kot 2001), which describes probabilistic changes in population sizes over time. In birth-death models, the probability of a birth or a death in a small period of time increases with the population size (Renshaw 1991). When the population size is small in a birth-death model, it is possible for extinction to occur due to a chance preponderance of deaths over births, even if the per-capita birth rate exceeds the per-capita death rate. Birth-death models are thus well suited to describe the demographic stochasticity that underlies replicative drift.

In our within-host birth-death model, pathogen extinction is equivalent to the clearance of the infection by the host. If the pathogen does not go extinct, its population eventually becomes large enough that the effects of stochasticity are negligible (Saaty 1961), leading to host death when the population reaches an upper threshold. In previous work we showed that linear birth-death models are insufficient to explain data on the speed of kill of the gypsy moth baculovirus, whereas models that allow for nonlinearities due to the immune system produce a better explanation for the data (Kennedy et al. 2014).

Our within-host model thus describes virus removal as the outcome of a process that begins with the insect’s immune system releasing chemicals that active the phenol-oxidase pathway. This release causes virus particles to be encapsulated and destroyed by host immune cells, and it also incapacitates the immune cell (Ashida and Brey 1998; Trudeau et al. 2001). Our model thus follows standard predator-prey-type immune-system models (Alizon and van Baalen 2008), in which the pathogen is the prey, and the immune cells are the predator, except that here the immune cells do not reproduce over the timescale of a single infection. The pathogen population in the model may then be driven to zero because of interactions with the host immune system, or it may persist long enough to overwhelm the host immune system, leading to exponential pathogen growth and eventual host death. Which outcome occurs depends on the initial pathogen population size and on demographic stochasticity during the infection.

In our model, the initial pathogen population size within a host is drawn from a Poisson distribution (fig. 1, Kennedy et al. 2014). If the infecting cadaver is composed of multiple strains, the model draws an initial population size for each strain from a multinomial distribution, such that the probability of sampling a particular strain from the infecting cadaver depends on the frequency of that strain in the cadaver. This process creates a transmission bottleneck. Next, the model tracks the population size of each virus strain over the course of the infection. Changes in the relative frequencies of these strains over time creates replicative drift. The host dies when the total pathogen population size exceeds an upper threshold. The frequency of virus strains at the time of host death determines the frequency of strains in the newly generated cadaver.

We model pathogen dynamics at the scale of the entire host population first by using a stochastic Susceptible-Exposed-Infected-Removed or “SEIR” model to describe epizootics (in our case the infected *I* class consists of infectious cadavers in the environment, which we symbolize as *P* for pathogen). Our SEIR model is modified to allow hosts to vary in infection risk, an important feature of gypsy moth virus transmission (Dwyer et al. 1997; Elderd et al. 2008), and to allow exposed hosts to be re-exposed, because infected gypsy moth larvae continue to consume foliage until shortly before death (Eakin et al. 2014). For computational convenience (Wearing et al. 2005), most SEIR models assume that the time between exposure and infectiousness follows a gamma distribution (Keeling and Rohani 2008). We instead allow this time to be determined by our within-host model, so that the within-host model is nested inside the stochastic SEIR model. As in the within-host model, the frequency of different virus strains at the population scale can drift due to chance events, such as the exposure of hosts to one cadaver and not another. Our between-host model therefore adds an additional source of drift to our nested models.

Over longer time scales, gypsy moth populations go through host-pathogen population cycles, in which host outbreaks are terminated by baculovirus epizootics. This pattern is typical of many forest defoliating insects (Moreau and Lucarotti 2007). The resulting predator-prey-type oscillations drive gypsy moth outbreaks at intervals of 5-9 years (Dwyer et al. 2004, we neglect the effects of the gypsy moth fungal pathogen *Entomophaga maimaiga*, which was having only modest effects in our study areas in Michigan, USA, when we collected our samples). Between insect outbreaks, virus infection rates are very low (Elderd et al. 2008), which may strengthen the effects of genetic drift.

Gypsy moths have only one generation per year, and therefore only one epizootic per year. We thus nest our within-host/SEIR-type model into a model that describes host reproduction and virus survival after the epizootic (fig. 1). The SEIR model determines which hosts die during the epizootic and which virus strains killed those hosts. This information is used in difference equations that describe the reproduction of the surviving hosts, the survival of the pathogen over the winter, and the evolution of host resistance, an important factor in gypsy moth outbreak cycles (Dwyer et al. 2000; Elderd et al. 2008).

By explicitly tracking the dynamics of individual hosts and pathogens, our model inherently includes the effects of genetic drift. We also tested whether a simple model of purifying selection, or a model of *de novo* mutation could explain the patterns of diversity in the data, without invoking drift within hosts. If these models were to fail to explain the patterns of diversity seen in our data, more complex models of evolution would need to be considered. In the gypsy moth baculovirus system, however, mutation rates are likely low (Rohrmann 2008; Sanjuán and Domingo-Calap 2016), spatial structure appears to be weak (Fujita 2007, Supplemental Information A), and evidence of selection acting within hosts is lacking (Supplemental Information H). Drift therefore seems likely to play a strong role in shaping virus diversity.

To show that the different sources of drift in our model are actually necessary to explain the data, we created three alternative models. All three alternative models simply the effects of genetic drift, but one also allows for effects of purifying selection. For the first alternative model, we simplified the effects of genetic drift by assuming that the relative frequencies of different virus strains within hosts do not change during pathogen population growth within hosts. To do this, we altered the model output such that the relative frequencies of virus strains released from a host upon host death were equal to the relative frequencies of virus strains just after the transmission bottleneck, thereby eliminating the effects of replicative drift (Fig. 1). For the second alternative model, we further simplified the effects of drift by assuming that the relative frequencies of virus strains at the end of an infection were the same as their relative frequencies in the infectious cadaver that initiated the infection, thereby eliminating both replicative drift and transmission bottlenecks (fig. 1). For the third alternative model, we added purifying selection to the second alternative model, which lacked both replicative drift and transmission bottlenecks. We did this by assuming that each host was susceptible to only a random subset of virus strains, so that exposure would only result in death if a host was susceptible to one or more virus strains in the cadaver to which it was exposed. The relative frequencies of virus strains released upon death was then equal to the relative frequencies of virus strains to which that host was susceptible to in the infecting cadaver.

### Baculovirus sequencing

We collected larvae from outbreaking gypsy moth populations in Michigan between 2000 and 2003 (Supplemental Information A), and we reared the larvae until they pupated or died of infection (Woods and Elkinton 1987). The virus population from each virus-killed larva was passaged once by infecting 75 larvae with liquefied cadaver to generate enough virus for DNA extraction. We then extracted DNA following a standard baculovirus DNA extraction protocol, and we amplified the DNA using whole genome amplification (REPLI-g UltraFast Mini kit from Qiagen).

We constructed sequencing libraries using the Nextera DNA Sample Prep Kit (Illumina-compatible, #GA0911-96) with custom barcodes to distinguish between the virus communities of different hosts. Our barcodes consisted of the first 96 indexes proposed by Meyer and Kircher (2010) (Supplemental Information A). Sequencing was carried out as two sets of libraries, run on individual lanes of a HiSeq2000 at the University of Illinois at Urbana-Champaign, producing 100 cycle single-end reads. Samples were separated by barcode using the standard Illumina pipeline, and adaptor contamination was removed using ‘trim galore’. Reads were mapped to the first sequenced gypsy moth baculovirus genome (Kuzio et al. 1999) using ‘bowtie2’ (Langmead and Salzberg 2012) with parameter set ‘very-fast’ (Supplemental Information A). Overall mean coverage was 886x, and varied across samples from 202x to 1497x (fig. S3). Variant calling was carried out using ‘VarScan’ version 2.3.9 (Koboldt et al. 2012). More details can be found in Supplemental Information A.

## Data and code availability

Sequence data generated in this study are available through the Sequence Read Archive (SRA) of the National Center for Biotechnology Information (NCBI) under BioProject ID PRJNA386565. Author-generated code is available at GitHub repository: https://github.com/dkenned1/KennedyDwyer.

## Acknowledgements

Our work was supported by NIH grant R01-GM096655 to G Dwyer, V Dukic, and B Rehill. Computational support was provided by the Computing Research Institute and the Research Computing Center at the University of Chicago.

